# Feedback information sharing in the human brain reflects bistable perception in the absence of report

**DOI:** 10.1101/2021.11.02.466729

**Authors:** Andres Canales-Johnson, Lola Beerendonk, Srivas Chennu, Matthew J. Davidson, Robin A.A. Ince, Simon van Gaal

## Abstract

In the search for the neural basis of conscious experience, perception and the cognitive processes associated with reporting perception are typically confounded as neural activity is recorded while participants explicitly report what they experience. Here we present a novel way to disentangle perception from report using eye-movement analysis techniques based on convolutional neural networks and neurodynamical analyses based on information theory. We use a bistable visual stimulus that instantiates two well-known properties of conscious perception: integration and differentiation. At any given moment, observers either perceive the stimulus as one integrated unitary object or as two differentiated objects that are clearly distinct from each other. Using electroencephalography, we show that measures of integration and differentiation based on information theory closely follow participants’ perceptual experience of those contents when switches were reported. We observed increased information integration between anterior to posterior electrodes (front to back) prior to a switch to the integrated percept, and higher information differentiation of anterior signals leading up to reporting the differentiated percept. Crucially, information integration was closely linked to perception and even observed in a no-report condition when perceptual transitions were inferred from eye movements alone. In contrast, the link between neural differentiation and perception was observed solely in the active report condition. Our results, therefore, suggest that perception and the processes associated with report require distinct amounts of anterior-posterior network communication and anterior information differentiation. While front-to-back directed information is associated with changes in the content of perception when viewing bistable visual stimuli, regardless of report, frontal information differentiation was absent in the no-report condition and therefore is not directly linked to perception *per se*.

## INTRODUCTION

Consciousness is subjective experience, the ‘what it is likeness’ of our experience, for example, when we perceive a certain scene or endure pain. Having such experiences may be the main reason why life matters to us, and it may set us apart from other smart but non-living “things”, such as your phone or the internet (1). Conscious experience is a truly subjective and private phenomenon and it cannot be observed directly from the outside. To objectively study and understand it, we have to get access to the inner subjective experience of others, for instance via self-report tasks or via behavioral tasks that we believe can capture conscious content (2). In this way, when combined with neuroimaging tools, the neural correlates of consciousness (NCCs) are sought (3). However, while doing so, what we observe in measures of brain activity may not be as pure as we hoped for. Our measurements of ‘consciousness’ may be cofounded with several cognitive factors, e.g., the act of reporting, attention, surprise, or decision-making, arising after conscious experience has emerged. This severely complicates our attempt to isolate the neural basis of conscious experience (4; 5; 6; 7; 8; 9). Here we address this complication, by assessing the influence of report and no-report task instructions on neural measures reflecting perceptual transitions in situations in which sensory input is ambiguous.

Perceptual ambiguity is a key phenomenon to study the brain mechanisms of conscious perception (10), often experimentally elicited by using ‘multistable stimuli’ (11). Multistability can be induced in several ways or using several ambiguous stimuli, for instance using binocular rivalry, structure from motion, the Necker cube, and motion-induced blindness. The common feature of such paradigms is that an ambiguous stimulus can be interpreted in two, or multiple ways, without changing the sensory input that reaches the senses. Confronted with this ambiguity, observers experience spontaneous fluctuations between interpretations of the stimulus.

Experimental evidence has varied regarding whether changes in perception when viewing multistable stimuli correlate with changes in early sensory or higher-order cortical activity. In support of higher loci, single-cell recordings in monkeys have revealed that the strongest perceptual modulations of neuronal firing occur in higher association cortices, including inferotemporal cortex (ITC) and dorsolateral prefrontal cortex (DLPFC) (12; 13; 14; 15). However, previous human fMRI studies have associated both early and higher sensory regions with perceptual transitions (16). For example, when face and house stimuli compete for perceptual dominance, category-specific regions in ITC activate more strongly for the dominant percept, even before observers indicate a perceptual switch via button press. Another common observation in human fMRI studies is the association of a large network of parietal and frontal brain areas, traditionally associated with attentional and cognitive functions, during the report of perceptual transitions (for an overview see (11)). One central and unresolved issue in consciousness science in general (8)), and multistable perception in particular, is what processes these large clusters of activations in the frontoparietal cortex reflect. The feedback account states that frontal regions actively exert a top-down influence on sensory brain regions to resolve perceptual ambiguity. Evidence in favor of this account shows that targeting specific nodes in this network using transcranial magnetic stimulation (TMS) can shape the rate of perceptual transitions, suggesting their causal influence in resolving, or even driving, perceptual ambiguity (11; 17; 18). The opposite feedforward account links frontal activity to processing of the consequences of perceiving, and thus processes occurring after perceptual ambiguity was resolved by posterior brain regions (19; 20; 18). Here, we contribute to this debate by assessing the extent to which feedforward and feedback processes correlate with changes in multistable perception, in the presence and absence of explicit report about the dominant percept.

By studying conscious contents in the presence and absence of explicit report during visual bistable perception, we here investigate the neural mechanisms underlying perceptual transitions, while dissociating perception from report, or other related factors. The bistable stimulus we use consists of two overlapping gratings, known as ambiguous plaids or moving plaids (21; 22). This ambiguous stimulus can be perceived as one plaid moving coherently in a vertical direction, or two plaids sliding across one another horizontally. The plaid stimulus was thus perceived as perceptually integrated (one object) or perceptually differentiated (two objects), respectively. Due to the unique direction of motion associated with these stimuli, this stimulus allows us to track perception in the absence of report by capitalizing on the occurrence of optokinetic nystagmus (OKN) (23; 24; 25; 26). OKN is a visually induced reflex, comprised of a combination of slow-phase and fast-phase eye movements that allow the eyes to follow objects in motion, for instance when looking at the trees alongside the road whilst moving past them in a car. Importantly, OKN follows perception when viewing bistable visual stimuli, which makes it useful for assessing perception in the absence of reports (23; 25; 26).

We focused on neural metrics inspired by information theory (27), to assess the neural underpinnings of awareness in report and no report conditions while viewing this integrated or differentiated percept. In a recent report-based study on perceptual ambiguity, it was shown that frontoparietal information integration, computed as the amount of information sharing between frontal and parietal EEG/ECoG signals, increased when participants reported an ambiguous auditory stimulus as perceptually integrated compared to when it was reported as perceptually differentiated. On the contrary, information differentiation, computed as the amount of information diversity within frontal or parietal EEG/ECoG signals separately, showed the opposite pattern within the same frontoparietal electrodes: it increased when participants reported the bistable stimulus as perceptually differentiated compared to perceptually integrated. This suggests that information integration and information differentiation go hand in hand with observers’ phenomenology of an integrated or differentiated percept of an ambiguous stimulus, a hypothesis that we explore in the visual modality here. One crucial open question so far, however, is whether the observed changes in neural integration and differentiation are dependent upon reporting or not because, in this previous study, observers had to explicitly report the perceptual switches by pressing buttons. We address this issue here by specifically relating neural metrics of integration and differentiation to changing percepts in both report and no-report conditions (5; 9). Further, we here also specify the directionality of neural information flow to relate feedforward and feedback activity to changes in perceptual experience, independent of the necessity to report.

## RESULTS

### Oculomotor signals

There were two experimental conditions that were performed in alternating runs. During the report runs (Figure 1A), observers pressed one button when the percept changed from vertical movement to horizontal movement and another button to indicate changes from horizontal movement to vertical movement (buttons were counterbalanced across blocks). During the no-report runs (Figure 1B), observers were instructed to remain central fixation and just passively view the stimulus, and therefore perceptual transitions were rendered task-irrelevant. Participants were not informed about the relationship between perception and oculomotor signals and not about our goal to infer perception from their eye signals.

**Figure 1:**
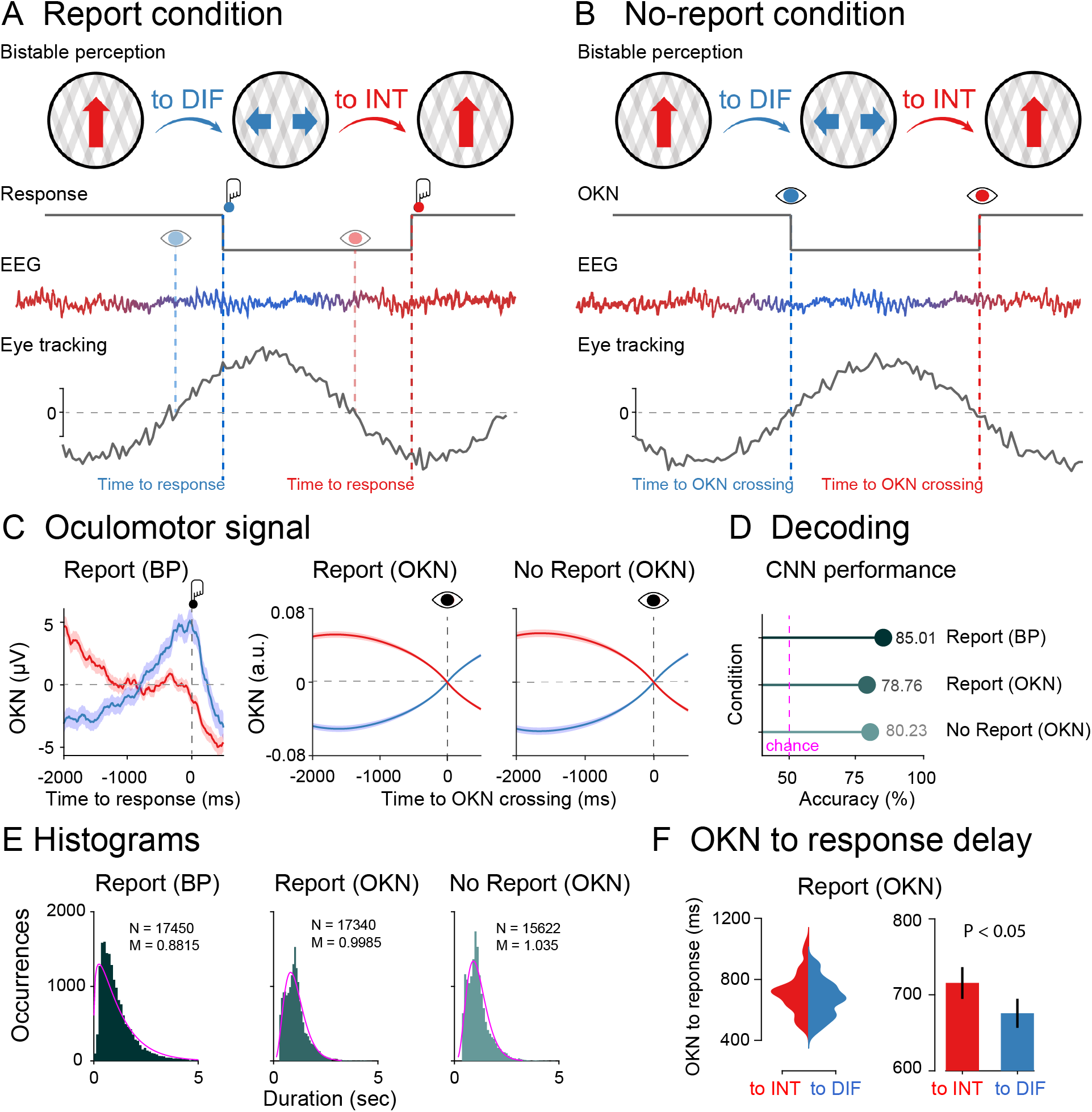
Experimental design, oculomotor decoding, and behavior. **(A)** Phenomenology during visual bistability in the report condition. Participants observed an ambiguous stimulus (moving plaids) that are experienced either as one plaid moving vertically (integrated percept; red arrow) or as two plaids moving horizontally (differentiated percept; blue arrows). Perceptual transitions occur either in integrated to differentiated direction (to DIF; blue) or in the differentiated to integrated direction (to INT; red). Middle row: Behavioral responses during the task. Participants pressed one button when perceiving that the integrated percept had fully changed into the differentiated percept (red button) and another button when perceiving that the differentiated percept had fully changed into the integrated percept (blue button). Bottom row: Dynamical analyses for EEG and oculomotor (eye tracking) signals. From the oculomotor response, we estimated the speed of the slow phase of the OKN (grey line) to infer participant’s perceptual content around OKN zero-crossings (to INT: red eye; to DIFF: blue eye) which coincides with their reports indexed by button presses (to INT: red button; to DIFF: blue button). **(B)** In the no-report condition observers passively viewed the bistable stimulus (no button presses) and perceptual alternations were inferred from oculomotor signals (to INT: red eye; to DIFF: blue eye). EEG y-axis represents voltage (microvolts) and x-axis time (milliseconds); OKN y-axis represents velocity (arbitrary units) and x-axis time (milliseconds). **(C)** Left panel: Raw oculomotor signal (in microvolts) locked to the button press (Report BP). Middle and Right panels: slow phase of the OKN (in arbitrary units) extracted from the oculomotor signal and locked to the OKN zero crossings (Report: OKN; No report: OKN). The slow phase of the OKN was obtained by computing the instantaneous velocity of the oculomotor signal smoothed over time (see Methods). Decoding accuracies **(D)** and histograms **(E)** for perceptual switches in the report condition locked to the button response, in the report condition locked to the OKN crossings, and in the no-report condition locked to the OKN crossings. **(F)** Delay between OKN crossings and button responses in the report condition (left panel: single-subject distributions; right panel: group-level analysis). Data is available at https://osf.io/a2f3v/.

We first characterized changes in oculomotor signals in both directions of perceptual change (vertical/horizontal) in the report condition time-locked to the button response. We computed the slow phase of the OKN (see Methods) as it distinguishes between visual percepts during binocular rivalry (25; 28; 29; 9; 30). OKN is particularly useful as positive and negative zero-crossings tend to precede the perceptual changes indicated by button presses. We observed OKN zero crossing going from positive to negative before the visual stimulus was reported as integrated, and zero crossing going from negative to positive, when the stimulus was reported as differentiated (Figure 1C). In the left panel of Figure 1C (Report: BP) we depicted the raw OKN time series time-locked to the button press (−200 to 500 ms) in microvolts. Next, for both the report (Report: OKN; Figure 1C) and no-report (No report: OKN; Figure 1C) conditions, after computing the OKN crossings in the continuous oculomotor signal, we labeled the data with their corresponding button press labels, and created epochs locked to the OKN crossings, which as expected, revealed a clear moment for the perceptual transitions (Figure 1C). In the middle and right panels of Figure 1C we plotted the slow phase of the OKN signal locked to zero-crossing in arbitrary units, which was computed as described in the Methods. It is important to note the similarity of OKN data in both report and no-report conditions, demonstrating its potential to capture the contents of perception in both cases.

To quantify the predictive value of the OKN crossings for identifying perceptual contents, we performed a decoding analysis using a convolutional neural network (CNN). To measure the generalization performance of the CNN, we split the OKN epochs and their corresponding button press labels into 3 parts: training, validation, and testing. 70% of the entire dataset was allocated to the training set. The remaining 30% were further split equally to create validation and test datasets, each containing 15% of the original data and labels (see Methods for details).

First, we trained a CNN to decode the button presses from the oculomotor signal using a cross-classification procedure in the report condition. We obtained a classification accuracy of 85% (Figure 1D). Next, in the same report condition, a second CNN was trained to decode the labels based on the OKN crossings, reaching a classification accuracy of 79%, indicating that OKN changes could reliably decode changes in perception. We finally estimated OKN crossings in the no-report data, labeled each crossing accordingly, and trained a third CNN to decode the labels, obtaining a classification accuracy of 80% (Figure 1D). Finally, another indication that our CNN accurately marks the occurrence of perceptual switches is that the number of switches in the report condition strongly correlates when based on button presses versus CNN performance (across subject Pearson’s r=0.88; p<0.001, see (23), for a similar analysis).

### Histograms

We next characterized the distributions of perceptual switches in the report and no-report condition, locked to button presses and OKN crossings (Figure 1E). All distributions were approximated well with a gamma distribution (p<0.001), a common observation in report-based rivalry paradigms (31). That the no-report dominance durations are well approximated by a typical gamma function further indicates that the variability (and phenomenology) in perceptual switches during the passive viewing condition is similar to active report conditions (see also (23; 31).

To investigate whether perceptual ambiguity is resolved differently when the stimulus is reported as integrated or differentiated, we computed the time delay between OKN crossings and button presses in the report condition. We observed a longer delay between OKN crossing and the button press when the stimulus was reported as integrated versus differentiated (t_1,39_=2.23; p=0.032; Cohen’s d=0.352, (Figure 1F). This increase in reaction time after a change in percept has occurred (as confirmed via OKN) suggests an increase in cognitive demand when reporting an integrated percept (transitioning from two objects to one) than when reporting a differentiated percept (going from one to two objects). We return to these features in the time-frequency characteristics of perceptual switches.

Overall, it is important to note the similarity of OKN data in both report and no-report conditions, demonstrating its ability to capture the content of perception in both cases. We observed the classic gamma functions for the distribution of percepts in each case. As a result, comparing our measures of integration and differentiation when time-locked to OKN crossings presents a powerful opportunity to assess the impact of report-based paradigms on the neural correlates of perception.

### Report condition locked to button presses

We first investigated the neural dynamics of information integration when participants reported perceptual switches by pressing buttons. Here, information integration refers to the transformation of inputs into outputs through recurrent neural dynamics. These nonlinear transformations are essential for pattern extraction in neural networks and may lead to signal amplification and broadcasting (32). To this end, we computed a metric of information integration known as Directed Information (dir-INFO) which quantifies directional connectivity between neural signals (33; 34). Compared to traditional causality detection methods based on linear models (e.g., Granger causality), dir-INFO is a model-free measure and can detect both linear and nonlinear functional relationships between brain signals. We took advantage of previous work that made this measure statistically robust when applied to neural data (35; 36; 33; 37). dir-INFO quantifies functional connectivity by measuring the degree to which the past of a “sender signal” *X* (e.g., EEG traces of anterior electrodes) predicts the future of another “receiver signal” *Y* (e.g., EEG traces of posterior electrodes), conditional on the past of the receiver signal *Y*. Thus, if there is significant dir-INFO between EEG signal *X* at one time, and EEG signal *Y* at a later time, this shows that signal *X* contains information about the future signal *Y*. Conditioning out the past of signal *Y* ensures the delayed interaction is providing new information over and above that available in the past of signal *Y*. For all dir-INFO analyses, we tested multiple delays from 0 ms to 500 ms (in steps of 4 ms) between the sender and receiver signal, which allows us to investigate the characteristic time delay of directed information transfer during perceptual switches between anterior and posterior signals. For all analyses reported here, we lock data to perceptual switches (either marked by a button press or eye-movement analysis) and inspect the EEG dynamics leading up to this perceptual switch. We have excluded all trials in which the previous perceptual switch occurred less than 2 seconds before the switch of interest (at time 0).

In the case of the report condition, (Figure 2A) shows dir-INFO between frontal and parietal signals, in both directions, so in the feedback direction (anterior to posterior electrodes) and in the feedforward direction (posterior to anterior electrodes, Figure S1A for electrode location). We plot feedforward and feedback-directed information as a function of signal delay between the two electrode sets, both when the moving plaids are reported as integrated (to INT; red color) as well as differentiated (to DIF; blue color). Signal delays are plotted on the y-axis, and testing time before the button press is plotted on the x-axis. A cluster-based permutation test performed on the difference between dir-INFO leading up to the integrated percept versus differentiated percept shows two significant clusters (cluster p<0.01). One cluster was observed indicating an increase in dir-INFO for the switch to integrated condition compared to the differentiated condition, at ∼ 50 ms delay between the time-period 1050 to 900 ms prior to the button press. A second cluster indicated an increase in dir-INFO prior to a switch to the integrated condition ∼250 ms delay of approximately 1200 to 500 ms before the button press (Figure 2). Importantly, these significant clusters were observed in the feedback direction only. Note that there are no button presses in the time window of interest, due to our trial exclusion procedure.

**Figure 2:**
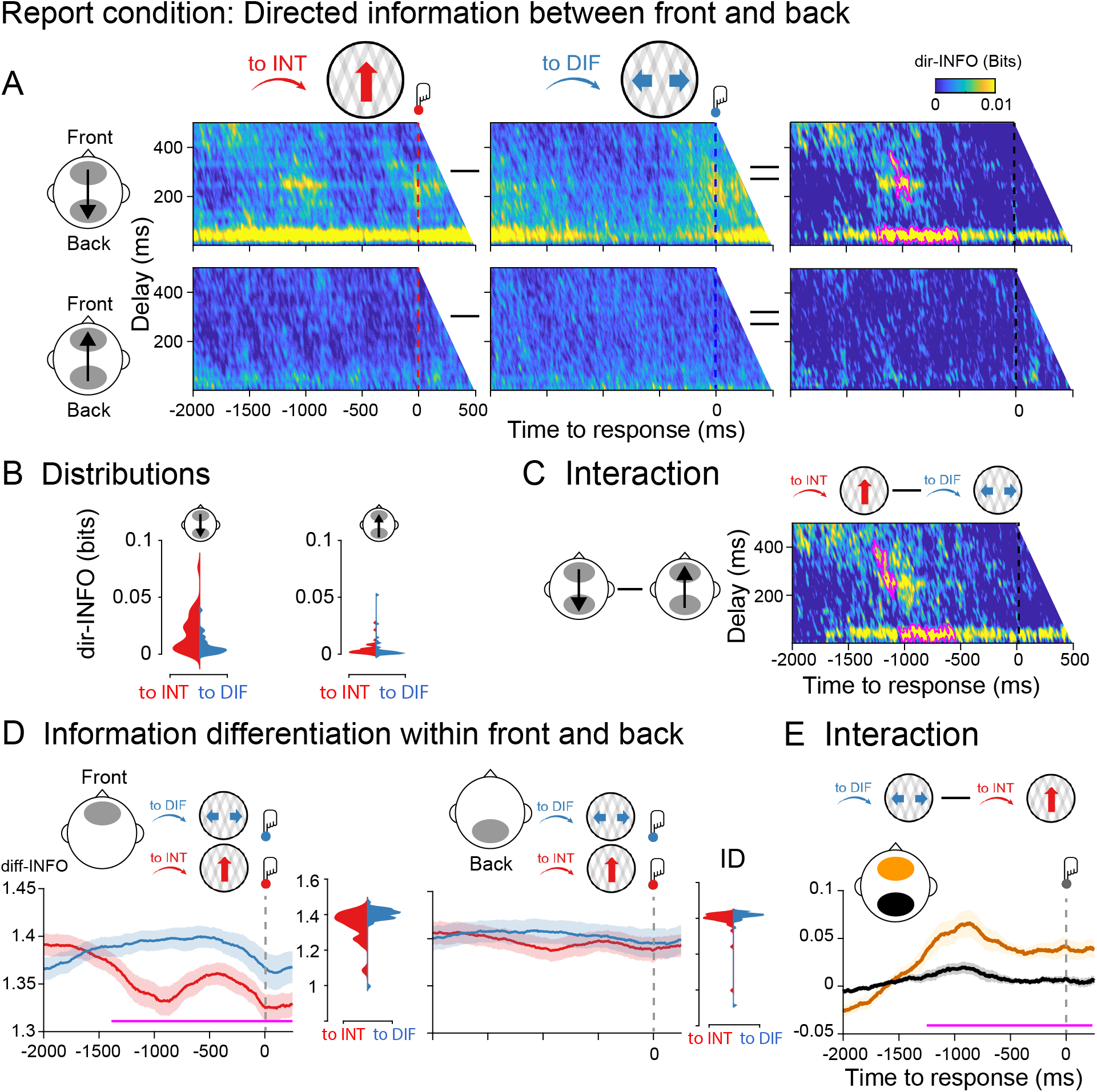
Information dynamics of the report condition locked to button responses. **A** Group-level dir-INFO between front and back ROIs (upper row) and between back and front ROIs (lower row) when moving plaids are reported as integrated (to INT; red color), reported as differentiated (to DIF; blue color), and the cluster-based permutation tests between the two. **B** Distribution of single-participant dir-INFO values for the frontal to the parietal direction (upper row) and for the parietal to the frontal direction (lower row). Values were extracted based on the significant clusters obtained in the frontal-to-parietal contrast (purple blobs). **C** Group-level dir-INFO statistical interaction effect computed as the difference between to INT and to DIFF trials between front and back ROIs (i.e., a double-subtraction). **D** Group-level diff-INFO within the frontal ROI and the corresponding single-subject diff-INFO values (left panel) when moving plaids are reported as differentiated and reported as integrated, and the same for the parietal ROI (right panel). **E** Group-level diff-INFO statistical interaction effect computed as the difference between to DIF and to INT trials between front (yellow color) and back (black color) ROIs. Data is available at https://osf.io/a2f3v/.

Next, we tested for an interaction between the direction of perceptual change (to integrated, to differentiated) and information direction (front to back, back to front). We performed a cluster-based permutation test on the difference between the perceptual switch (to INT minus to DIFF trials) and information direction (feedback minus feedforward). We observed two significant clusters (cluster p<0.01) showing stronger dir-INFO when perception switched to integrated as compared to differentiated in the front-to-back direction, but not in the back-to-front direction (Figure 2C). Finally, to test for the spatial specificity of the dir-INFO effect (i.e., the anterior-posterior direction), we computed dir-INFO in the right-left direction using temporal electrodes (Figure S1B). No dir-INFO differences between perceptual switches were observed in the right-to-left direction, nor in the left-to-right direction (Figure S2).

We next analyzed the dynamics of information differentiation (K-complexity) within frontal and parietal signals separately. Neural differentiation metrics quantify the diversity of information patterns within brain signals and it has been useful for distinguishing between conscious states (38; 39; 40) and conscious contents previously (27). We performed cluster-based permutation testing on the difference between the Information Differentiation (ID) time series leading up to the integrated percept versus the differentiated percept for the same set of anterior (front) and posterior (back) electrodes. As expected, significantly increased diff-INFO was observed before the bistable stimulus was reported as differentiated as compared to when was reported as integrated. This effect was observed at approximately -1300 to 250 ms around the button press indicating the upcoming perceptual switch (Figure 2E), note that in the 2000 ms before the perceptual switch indicated by the response no previous switches are incorporated). Importantly, no such effect was observed for the posterior electrodes (Figure 2D).

We next tested for an interaction between the direction of perceptual change and information differentiation by subtracting between to DIF and to INT trials within frontal and back ROIs (Figure 2E). As predicted, a cluster-based permutation test revealed an interaction between perceptual change, perceptual direction, and ROI based on information differentiation (p<0.01), showing higher diff-INFO in the anterior region prior to a change to seeing the differentiated percept, without changes to diff-INFO in the posterior regions (Figure 2E). Finally, we computed diff-INFO in the right and left ROI using the temporal electrodes. No diff-INFO differences between perceptual switches were observed in the right nor left ROI (Figure S3).

Taken together, and consistent with previously reported effects in the auditory domain (41), these results show that having an integrated versus a differentiated percept, goes hand in hand with front-to-back neural directed information integration and frontal differentiation brain measures. Next, we aim to establish to what extent these effects depend on the necessity for reporting perceptual transitions (task relevance of switches) and we further specify the timing of these effects by locking our analyses more closely to the perceptual transitions based on OKN crossings.

### Report condition locked to OKN crossings

The dir-INFO analyses were performed in the same way as in the report condition but this time locked to OKN crossings instead of button presses. We observed significant clusters of increased dir-INFO when the visual stimulus was perceived as integrated as compared to differentiated, again only in the feedback direction (Figure 3A).

**Figure 3:**
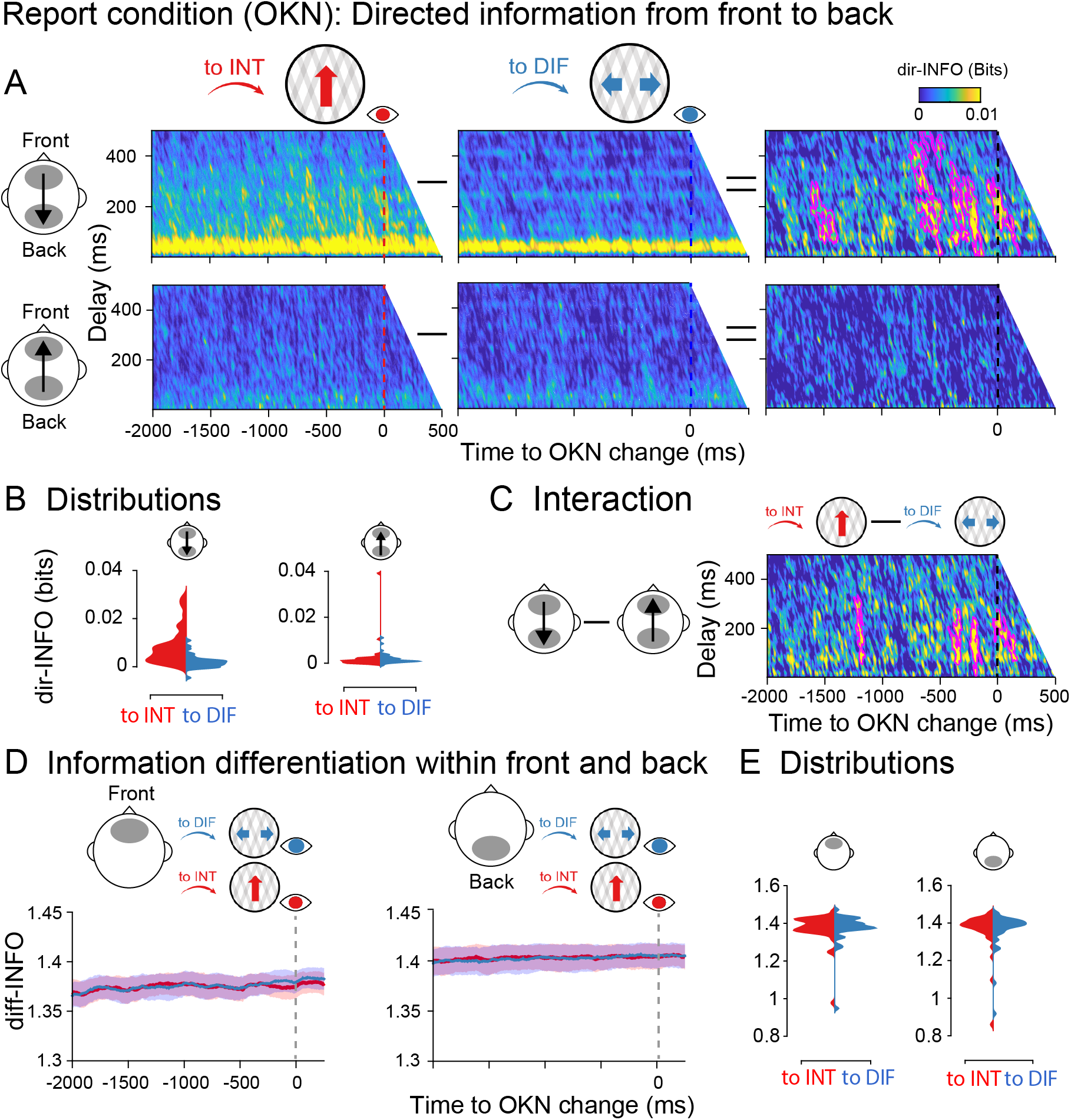
Information dynamics of the report condition locked to OKN crossings. Specifics between **A-D** are identical to Figure 2. **E** Single participant diff-INFO values were extracted using as a reference the significant time window observed in Figure 2D. Data is available at https://osf.io/a2f3v/.

Significant clusters spanned delays between 50 to 450 ms and occurred roughly -1600 to 100 ms around the moment OKN crossed. Interestingly, significant time points were observed much closer in time to the perceptual switches when time-locked to OKN crossings, than when locked to responses (Figure 2A). Similar interaction analysis between switch direction and integration direction (feedback vs feedforward) was performed. The difference in directed information between integrated and differentiated percepts was stronger for the feedback direction than the feedforward direction (interaction cluster p<0.01; Figure 3C).

For information differentiation however, locked to the OKN crossings in the report condition, no robust clusters were observed (Figure 3D), neither for the front (left panel) nor the back (right panel) ROIs. As a strong diff-INFO effect was observed in this same data when aligned to button-press (Figure 2D), we note that this measure may capture processes involved in (manual) report, rather than in the experience of a perceptual state. We return to this nuance in our Discussion.

### No report condition

After establishing the dynamics of information integration and differentiation in the report condition, and finding that differentiation measures, but not integration measures, were dependent on report, we next analyzed both during the no-report condition (i.e., locked to the OKN crossings). Similar to the report condition, a cluster-based permutation test revealed significant clusters of increased dir-INFO when the visual stimulus was perceived as integrated as compared to differentiated, but uniquely in the feedback direction (Figure 4A). The observed clusters showed a similar range of delays (around 50 to 400 ms) and they were observed in similar time windows (−600 to 0 ms around the OKN crossings) as in the report condition when time-locked to the OKN (Figure 3). We also observed significant clusters showing an interaction between perceptual switch and information direction (p<0.01), showing stronger integra-tion when the percept switched to integrated as compared to differentiated, and more so in the feedback than feedforward direction (Figure 4C).

**Figure 4:**
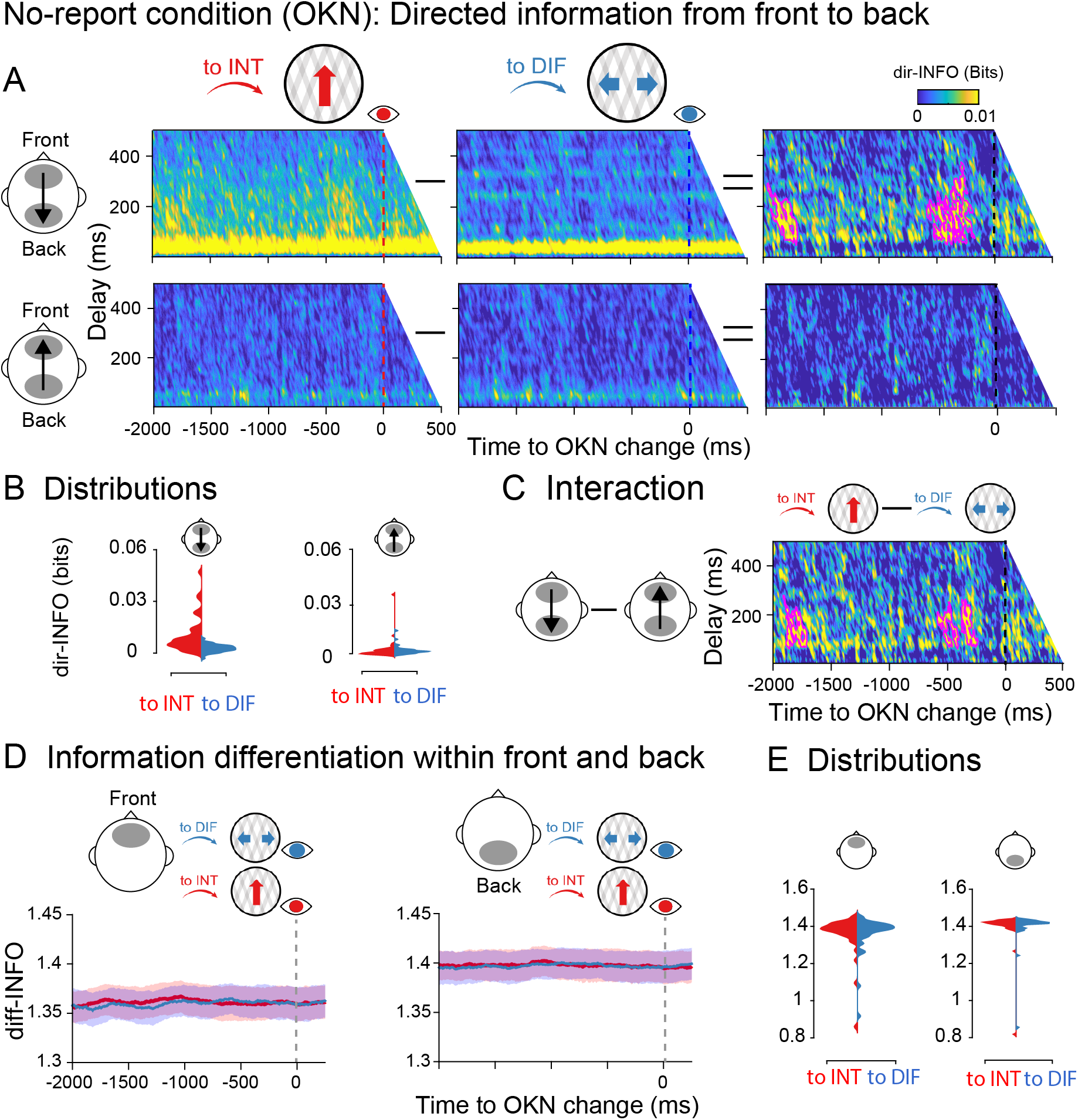
Information dynamics of the no-report condition locked to OKN crossings. Specifics between **A-D** are identical to Figure 2. **E** Single participant diff-INFO values were extracted using as a reference the significant time window observed in Figure 2D. Data is available at https://osf.io/a2f3v/.

Finally, we analyzed the dynamics of information differentiation during the no-report condition. Again, no robust diff-INFO temporal clusters were observed when testing the entire time window -2000 to 500 ms around the OKN crossing between perceptual switches neither in the front nor in the back ROIs (Figure 4D). Taken together, these results indicate that directed information distinguishes visual contents even in the absence of explicit report, while information differentiation in anterior locations is dependent upon report.

### Eye movements cannot explain the observed neural effects

We would like to note that the observed EEG effects are unlikely to be the result of differences in eye-movement signals between conditions for several reasons. First, measures of neural differentiation were specific to the report-locked analyses, although similar eye-movement patterns were present in all conditions. Second, the difference in the directionality of the dir-INFO effects (presence of a dir-INFO effect in the feedback but not feedforward direction, no effects for temporal electrodes) indicates that general eye-movement confounds are also not likely. Third, OKN signals gradually change over time over a period of 2 seconds leading up to the perceptual switch, with highly similar time courses for both types of percepts (see Figure 1C)). The neural effects we observed do not reflect such a gradual build-up and differ based on the perceived content which differs from what one would expect based on the pattern of the OKN signals. Fourth, it is also unlikely that eye-related signals (e.g. muscle activity) get somehow embedded in the EEG recordings and drive our anterior-to-posterior effects. The idea would be that if this signal is first measured on frontal and then on posterior electrodes, transfer entropy or source conduction would increase measures of directed information flow. This interpretation is unlikely given that we did not observe any differences in OKN signals when comparing to INT and to DIFF periods in the time window 1200 to 500 ms before the button press, where we observed the differences in our neural measures in the report condition (Figure 1C; dependent-samples t-test (to INT, to DIF): t_1,39_=0.093; p=0.926; BF_01_=5.77). Fifth, when the eyes change their direction maximally, that is at the zero crossings where the derivative of the slope is largest (Figure 1C), or when the eyes most strongly indicate a certain perceptual state (at the peak of the OKN signal in Figure 1C), dir-INFO in the feedback direction was not maximal. If the eyes drive the neural effects, a peak in dir-INFO would be expected.

Finally, and most importantly, we have re-analyzed a previously published dataset in which we used an auditory bistable stimulus in combination with EEG measurements (27). In that study, participants listened to a sequence of tones and indicated with a button press whether they experienced either a single stream (perceptual integration) or two parallel streams (perceptual differentiation) of sounds. In that dataset, there were no systematic associations between eye movements and perceptual switches due to the auditory nature of the task. In Figure S4 we report the dir-INFO analysis showing a highly similar pattern of results as reported here for bistable visual stimuli. Again, we observed increased dir-INFO leading up to integrated percepts, and uniquely in the feedback direction (see Figure S4 for details). Together, these considerations and additional results from auditory bistability indicate that the results observed here are unlikely to be caused by incidental differences in eye-movement patterns between conditions or perceptual states.

### Time-frequency characteristics of perceptual switches

To compare the integration and differentiation results with more traditional electrophysiological features of multistable perception, we computed the time-frequency profile of perceptual switches to the differentiated and integrated percepts in Figure 5 (using cluster-based corrections for multiple comparisons). We observed increased oscillatory power in a broad frequency band from 1 to 18 Hz at frontal electrode sites, preceding a perceptual transition to an integrated compared to a differentiated percept. This was only observed in the button press locked analyses of the report condition, not in any of the other conditions. Interestingly, the same pattern of results was observed in the frontal complexity measure (diff-INFO) (Figure 2D,E). Complexity measures have been linked to changes in frequency dynamics (42), and in particular, an increase in low-frequency power can increase the redundancy in neural signals, decreasing information complexity (43; 44; 45). Accordingly, we observed that in the time window that overlaps both measures (from -1050 ms before the response onset), the decrease in complexity negatively correlated with time-frequency power (Pearson’s r=-0.39; p=0.013). We return to this result in our Discussion.

**Figure 5:**
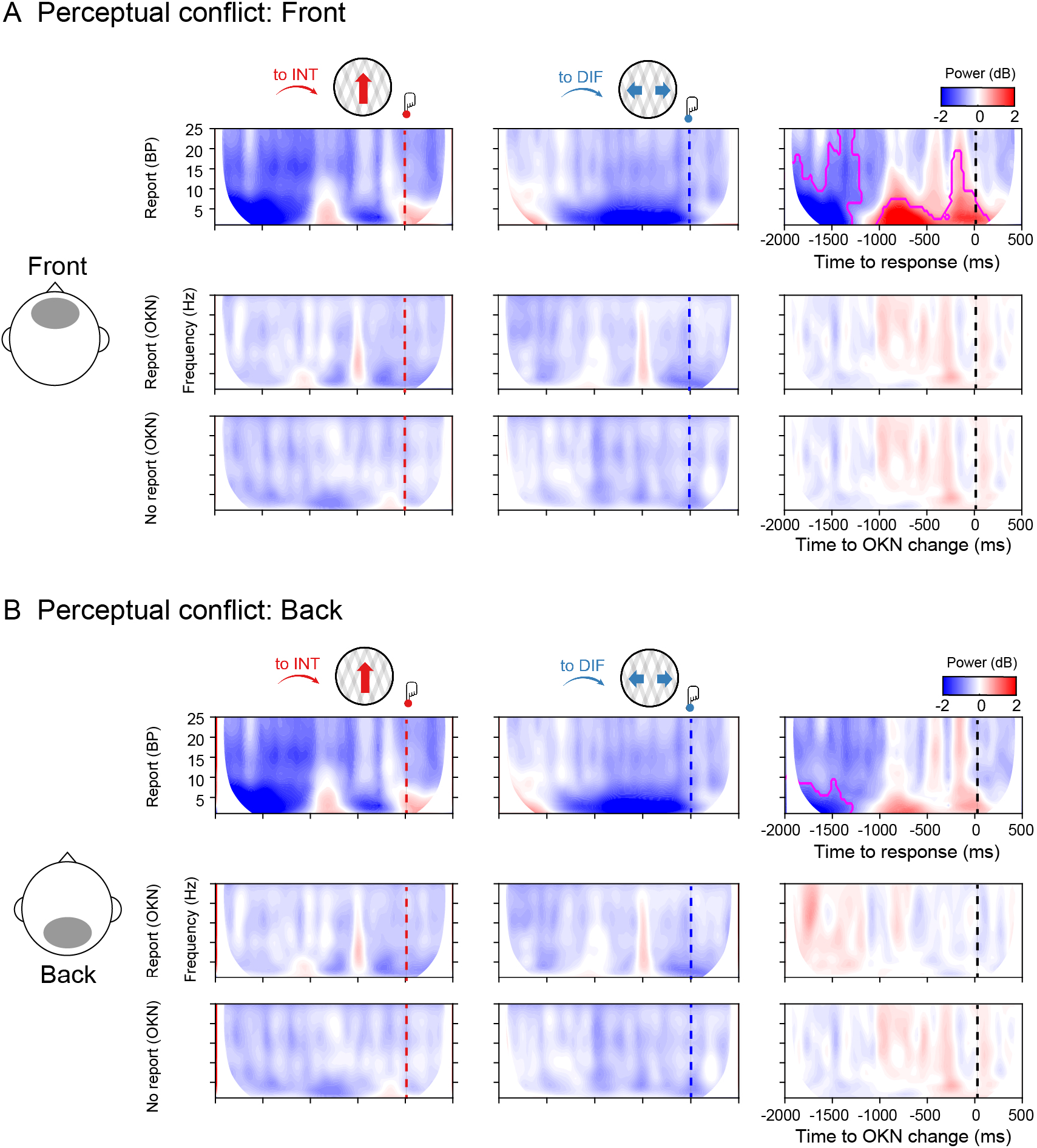
Spectral power in the report and no report conditions. **A** Group-level spectral power in the front ROI for the report condition (upper row), report condition locked to OKN crossings (middle row), and no report condition (lower row), when moving plaids are reported as integrated (to INT; red color), reported as differentiated (to DIF; blue color), and the cluster-based permutation tests between the two. **B** Same as **A** but for the back ROI. Data is available at https://osf.io/a2f3v/.

## DISCUSSION

Perceptual rivalry is the phenomenon that the perceptual interpretation of a bistable visual stimulus alternates over time in the absence of physical changes in the presented sensory input. The use of perceptual rivalry, as a tool, has provided fundamental insight into what neural processes may reflect the content of our conscious experience. However, although great progress has been made in unraveling neural processes underlying the competition between neural representations and the perceptual change between alternative percepts, it has proven notoriously difficult to separate neural correlates of perceptual switches, from processes that have another origin and are associated, for instance, with report, attention, surprise or commitment to a decision (8; 9). Here we aimed to tackle this issue using an experimental set-up that contained four crucial ingredients. First, we experimentally introduced conditions in which perceptual switches were task-relevant and had to be reported about, versus conditions in which perceptual transitions were irrelevant and bistable stimuli just had to be viewed passively (23; 25; 46). Second, we relied on neural measures with millisecond temporal resolution, allowing us to pinpoint the relevant neural processes as they evolve over time. Third, we capitalized on novel information-based measures that have been shown to reflect the phenomenology of conscious perception (33; 27), allowing us to track specific perceptual content over time, while it naturally alternates. Fourth and finally, we introduced a novel way to analyze eye-tracking data to be able to pinpoint when precisely in time perceptual interpretations alternate, allowing us to time-lock our analysis to focus on processing leading up to the perceptual change (times prior to OKN crossing), from processing involved in translating perception into action, as well as other cognitive confounds arising after the perceptual switch (times after OKN, prior to button-press). This effort has led to several novel results and conclusions, which are summarized below.

We showed that perceptual switches can systematically be inferred from eye-movement measurements during passive viewing of a bistable visual stimulus by capitalizing on OKN measures in combination with deep neural network modeling. This confirms previous work that has shown that OKN and pupil measures can act as reliable indicators of the dynamics of perceptual and binocular rivalry (25; 28; 26). In fact, percept durations varied commonly during experimental blocks in which perceptual switches had to be reported, and were therefore task-relevant, and blocks in which observers passively viewed the bistable stimulus (perceptual switches are task-irrelevant). In both conditions, the observer’s percept durations followed a right-skewed gamma distribution as has been observed previously across a wide range of bistability paradigms (31), suggesting that the phenomenology of perception was similar during active report and passive viewing. Next, we showed that when the perception of a bistable stimulus was explicitly reported, directed information from anterior to posterior signals was increased before the observers reported perceiving the integrated stimulus (one plaid moving upwards) compared to when the differentiated percept was reported. Note that although the posterior signals were not the source but the receiver of the information flow, posterior signals were still involved in the processing of the percept as the feedback pattern only emerged due to the statistical relationship between anterior and posterior signals. A different situation was observed when signals were analyzed in isolation using the information differentiation metric. Thus, before the percept was reported as differentiated (two stimuli, each moving sideways in a different direction), we observed higher information differentiation of anterior signals, as compared to integrated percepts, before the perceptual switch indicated by the button press. This effect was not observed on posterior sensors. These results indicate that our information-based measures derived from electrophysiological activity capture and track the phenomenology of the percept during perceptual ambiguity. These results are in agreement with our previous report-based study on auditory bistable perception showing a correspondence between perceptual integration and differentiation and neural metrics of information integration and differentiation (27). During report, when dir-INFO is high an integrated percept is likely to be perceived, whereas when diff-INFO is high, a differentiated percept is more likely to be perceived.

When perceptual changes were inferred from eye movements during passive viewing, we observed that our directed integration measure still tracked the perceptual state of the observer, but uniquely so in the feedback direction (from anterior to posterior signals). However, information differentiation measures no longer tracked the evolving percept, suggesting that our differentiation measures may reflect other processes not directly linked to perception, such as differences in task set or for example differences in attention due to the necessity to report. In sum, the relative strength of directed information in the feedback direction indicated which percept likely dominated during passive viewing, whereas neural differentiation does not. We would like to note that this means that directed information is thus a “symmetrical measure” in the sense that when it is high it reflects that an integrated percept is likely to be dominant, whereas when it is low, a differentiated stimulus is likely dominating perception.

During ambiguous perception (e.g. bistability, rivalry), feedback connections are thought to be critical for the comparison of internally generated predictions of sensory input with actual inputs (20; 18). Based on the neurobiology of feedback connections, they are vastly more numerous and divergent than feedforward ones, i.e. fewer neurons project in a feedforward manner, compared to a feedback one (47). We believe this dominance of feedback signals may account for the increase in directed information we observed prior to perceiving the integrated (vertical) motion percept (Figure 2A, 3A, 4A). More specifically, for our bidirectional plaid stimulus, perceiving an integrated percept of unified vertical motion required the holistic combination of local signals across the visual field. Thus, for perceptual integration, local signals from low-level areas may have required enhanced feedback modulations from higher areas to integrate visual information across space (48). This integration can be seen as hypothesis testing in which the high-level interpretation can inform the low-level features where feedback projections are considered to enhance neural activity (49; 50; 51). Supporting this view, recent studies on binocular rivalry have shown that PFC neurons increase their firing rate before the occurrence of perceptual switches both in monkeys (29) and humans (52). In the case of multiunit activity (MUA) recordings in humans, increased frontal activation precedes the activity of lower areas such as MT, suggesting that the frontal cortex biases sensory information in the lower areas towards one of the two perceptual contents in a top-down feedback manner (52). Thus, our results are in agreement with this prior research, as stronger top-down modulation preceded the integrated interpretation of our bistable plaid stimulus.

It is important to stress that we used information-specific EEG measures, which allowed us to track the contents of rivalrous states while quantifying the directionality of integration for our observers. This approach is similar to previous binocular rivalry studies using for instance alternating face/house stimuli and isolating (the competition between) neural activity in feature or category-selective brain regions, either using electrophysiology (20) or fMRI (16; 53). The use of these EEG information-based measures enabled us to track the neural signatures of the content of perception with high temporal precision before perception alternated, similar to influential electrophysiological studies using feature-specific neural responses (12; 54; 29; 52). Due to the low temporal resolution of the BOLD signal, to be able to isolate the neural cause of perceptual switches during rivalry, previous fMRI studies have often included a replay condition with yoked switches that are simulated using video replay. These replay trials mimic perceptual switches but have no neural cause as they are not driven by fluctuations in brain activity. fMRI studies have shown that the dorsolateral prefrontal cortex (dlPFC) activates stronger for perceptual switches during rivalry (as compared to baseline), and often also activates stronger during rivalry than during replay (31; 25; 7; 46; 55). Frässle and colleagues showed that when comparing rivalry to replay conditions, there was an increase in dlPFC activity when actively reporting on perceptual changes. However, the difference between rivalry and replay diminished in pre-frontal areas during passive viewing (25), suggesting that actively reporting on rivalry recruits additional prefrontal resources compared to reporting on replay, which is not required for passively experiencing a change in percept (see also e.g. (16)). Similarly, Brascamp and colleagues (31) have shown that prefrontal activation in fMRI (or in their words “responses in executive brain areas” in general) may be driven by the fact that perceptual switches often draw attention (possibly because they are surprising and relevant) and require report, but are not related to perceptual transitions per se. They have shown this by designing a task in which switches in perception between two stimuli (dots moving in different directions in the two eyes) could go unnoticed by observers, as if these switches were preconscious (potentially accessible but not actually accessed) (56). Although these switches remained unnoticed to the participants, sensory brain regions still showed neural activity pat-terns associated with those switches, whereas switch-related modulations in executive areas were minimized (although still numerically higher than baseline). Based on such results it has been argued that PFC involvement during rivalry may potentially be related to report, or other consequences of perceptual switches, but that the PFC does not drive those switches (25). Interestingly, our information differentiation (diff-INFO) results also suggest that frontal information differentiation may reflect processing associated with report rather than perception *per se*. In the report condition, we found that reporting the stimulus as integrated took longer than reporting it as differentiated, indicative of a task-dependent increase in the cognitive demand of disambiguating the stimulus identity. It may be that these differences can explain the frontal information differentiation effect observed during the report of perceptual switches, compared to the absence of this effect during the passive viewing condition. Knapen and colleagues (7) showed that large parts of the prefrontal network, again especially dorsolateral PFC, activate stronger perceptual transitions with longer durations, so when the system takes longer to settle in a particular perceptual state (the transition phase between interpretations is longer) than when the transition phase is shorter. Finally, a recent study (18) has shown that transcranial stimulation of the right inferior frontal cortex (IFC) reduced the occurrence of perceptual changes for bistable stimuli when they had to be actively reported (see also (57) for similar results while stimulating the DLPFC). The authors argued that the IFC may register perceptual conflicts (or the mismatch/prediction error signal) between two possible perceptual interpretations, gradually building up towards the perceptual switch. Therefore, the IFC may be influencing the competition between pools of neurons coding for the different percepts in the visual cortex, thereby “steering” or “co-determining” conscious perception (19). Our front-to-back dir-INFO results are more in line with this latter finding, suggesting that even during no report conditions, subtle influences from frontal signals co-determine perceptual switches during visual bistability (and report-based auditory bistability (Figure S4), but see (57)).

The experiment designed here can be regarded as a so-called “no-report paradigm” and not the stricter “no-cognition paradigm”, because in principle when one eliminates any type of explicit report, this does not necessarily remove all possible processes prior to reporting that could happen throughout the experiment (5). Observers may still reason, think or reflect about the presented bistable stimulus and perceptual alternations. This issue is similar to previous attempts to minimalize cognitive processes during perceptual rivalry (23; 25; 46). In our experiment, the crucial aspect is that perceptual switches were task-irrelevant during the passive viewing condition and task-relevant during the report condition (58). We cannot and did not control what observers were doing during passive viewing and therefore do not know whether and how observers were reasoning about the perceived stimulus or the occurrence of perceptual switches. Therefore, we only intended to arbitrate between task/report-related and perception-related information-based measures of perceptual transitions. Interestingly, using a similar set-up, combining OKN measures with manipulation of task-relevance of perceptual switches, it has recently been shown that also in the pupil response, task/report and perception-related dilations and constrictions can be separated (23). Finally, a cautionary note on the experimental design is that it is possible that participants were preparing an overt response in both conditions, but just withheld their response in the no-report condition. This may be a consequence of alternating blocks, in which we chose to equate the conditions as much as possible (e.g. match rates of perceptual learning, fatigue, and drowsiness over time between conditions). It is possible that perceptual changes were attended to in the no-report blocks, although they were not task-relevant.

We note, however, that we also observed frontal theta-band power increases uniquely in manual report conditions – supporting the separation between report and no-report states in our design. Frontal-central theta power has often been related to the processing of conflict, both at the level of stimuli as well as for conflicting stimulus-response mappings (59; 60). In the time-frequency response-locked analyses we observed increased theta-band power leading up to the integrated compared to differentiated percept, as well as longer reaction times between OKN crossings and report. This pattern of results was in agreement with the analysis of information differentiation in the report condition (Figure 2D), which was also only observed in the response-locked analyses. Combined, these results suggest that there may be additional differences between reporting an integrated and differentiated percept, that are contingent on subjective conflict, an exciting possibility for future research that may unify mechanisms of perceptual and cognitive decision-making (61).

Despite much debate in the field, influential theories of consciousness (62), such as Global Neuronal Workspace theory, Recurrent Processing theory, and Integrated Information Theory, agree that the common property of conscious experience relates to the brain’s capacity to integrate information through recurrent processing (the combination of lateral and feedback interactions), enhancing cortico-cortical interactions at a local scale (between nearby regions) or global scale (between distant regions) (63; 64; 65). Note that such consciousness theories are mostly concerned with explaining which neural processes can be observed when sensory information “crosses the threshold” from subliminal (unconscious) to phenomenal or access consciousness, and thus aim to isolate the necessary ingredients constituent to conscious experience. Instead, we focus here on which neural processes influence the competition between alternative interpretations of ambiguous visual input (see (19)) for a short review of this difference). In light of that latter debate, we show that the amount of directed information flow between anterior to posterior electrodes reflects the likelihood of perceiving the integrated version of an ambiguous stimulus during perceptual ambiguity, even in a context where report is not required. The low spatial resolution of our EEG measurements, however, precludes any claims about the specific origins of these observed feedback signals and future studies are needed to obtain more spatial specificity as well as a more mechanistic explanation of the neural processes the directed information measure reflects.

In conclusion, our results suggest that the relevant dynamical mechanism for perceiving different contents during visual bistability, controlling for many factors associated with reporting perception, is the directed information between frontal and posterior signals, rather than the isolated information differentiation contained within the front or the back of the brain.

## Funding

This research was supported by an ABC Talent Grant of the University of Amsterdam (ACJ, SVG), a grant from the H2020 European Research Council (ERC STG 715605, SVG), and a BIAL Foundation Grant 2020/2021 (ID: A-29477, SVG, ACJ).

## Authorship contributions

Conceived and designed the experiments: ACJ, SVG. Performed the experiments: ACJ, LB. Analyzed the data: ACJ, MD. Contributed reagents/materials/analysis tools: RI, SC. Wrote the paper: ACJ, LB, SC, RI, MD, SVG.

## Acknowledgments

We thank Dr. William J. Harrison, Dr. Maartje de Jong, Dr. William Gilpin, Dr. Martin Vinck, and Dr. Tomas Knapen for contributing to valuable discussions and insights and proofreading the manuscript. This manuscript is dedicated to the memory of Prof. Walter J. Freeman (1927 - 2016) whose pioneering work on Neurodynamics has inspired and ignited countless meaningful insights during the execution of this project.

## Conflict of Interest

None declared.

## METHODS

### Participants

Forty-two participants (31 females, 2 left-handed) aged between 18 and 35 (M = 20 years, SD = 3.73), recruited from the University of Amsterdam (Amsterdam, the Netherlands) participated in this study for monetary compensation. All participants had a normal or corrected-to-normal vision. The study was approved by the institutional review board (IRB) of the Psychology department of the University of Amsterdam (project ID: BC-8686), and written informed consent was obtained from all participants after the explanation of the experimental protocol. One participant was excluded during data collection for not following the task instructions. Another participant was excluded during the analyses because of very strong artifacts in the EEG data.

### Stimulus

The stimulus consisted of two overlapping semi-transparent half-wave gratings, known as ambiguous plaids or moving plaids (66; 67; 22). Each grating was a sinusoid (0.33 cycles per degree of visual angle), clipped to include only positive contrast luminance, positioned at the center of an aperture (diameter = 36.03). Peak contrast was set to 0.025 relative to the uniform grey background. The encoding gamma was set to 2. The orientation of each grating was 15° and -15°. Motion was created by changing the phase of the grating on each frame (6 Hz motion). We selected this specific stimulus for three reasons. Firstly, the stimulus is ambiguous, meaning that perception alternates whilst the sensory input remains constant. This approach is considered to be one of the most powerful methodologies to study the neural underpinnings of phenomenal consciousness (9). Secondly, the stimulus can be perceived as one grating moving coherently or two gratings sliding across one another. In other words, gratings are either perceptually integrated or perceptually differentiated, respectively. Note that the ambiguous plaids stimulus is considered to be tristable (rather than bistable) because it has one integrated percept (the gratings moving together as a single pattern) and two differentiated percepts (the gratings sliding across one another) with alternating depth order (which grating is perceived as foreground and which as background) (66). However, we treat the stimulus as if it were bistable because we instruct participants to respond to changes from the integrated percept to the differentiated percept and vice versa. The last reason for using this stimulus is that it allows us to track perception in the absence of responses by exploiting the occurrence of optokinetic nystagmus (OKN) (68; 25; 26).

### Experimental design and procedure

There were two experimental conditions (i.e. report and no-report) that were performed in alternating runs of four minutes each (Figure 1A). Participants performed twenty runs in total, split up into five blocks consisting of two runs of each condition. The order within the blocks was always the same and started with a no-report run (i.e. no-report, report, no-report, report). Note that the same stimulus was used for both conditions and that the stimulus was constant throughout each run. Only the direction of motion of the stimulus (i.e. upwards or downwards) was changed every two runs in order to exclude the direction of the stimulus as a confounding factor (Figure 1A). During the report runs, participants were instructed to press one button when the percept changed from vertical movement to horizontal movement and another button to indicate changes from horizontal movement to vertical movement (Figure 1B,C) (i.e. 2AFC). The buttons were operated with the index fingers of both hands and the contingencies of the buttons were counterbalanced across participants. During the no-report runs, participants were instructed to remain focused on the stimulus at all times and to relax their hands on the desk in front of them (i.e. passive viewing of the stimulus). Throughout the experiment, participants remained unaware of the relationship between perception and eye movements, and they were not informed about the influence of OKN on their perceptual state.

The experiment was programmed and executed using the Psychophysics Toolbox (version 3.0.14; Brainard, 1997) and Eyelink Toolbox extensions (Cornelissen, Peters, and Palmer, 2002) for MATLAB (R2016a, MathWorks, Inc., Natick, MA, US). Stimuli were presented on an Asus VG236H LCD monitor (23” diagonal, 1920 × 1080 pixel resolution; 120 Hz refresh rate) at a viewing distance of 50 cm.

### Electroencephalography (EEG) recording and preprocessing

EEG signals were acquired through a 64-channel Biosemi ActiveTwo system (Biosemi, Amsterdam, the Netherlands) with two online references (CMS and DRL) placed according to the international 10-20 system. Two external electrodes were placed on the earlobes for possible use as offline references. Four additional external electrodes were used to record vertical and horizontal eye movements, adding up to 72 channels in total. Data were sampled at 512 Hz. Preprocessing was done by means of custom-made MATLAB (R2016a, The MathWorks, Inc.) scripts supported by EEGLAB (69). Continuous EEG data were first down-sampled to 250 Hz and filtered between 1-100 Hz. Data from the 64 channels over the scalp surface (i.e. reference electrodes and external electrodes excluded) were retained for further analyses. All twenty runs were extracted and subsequently appended in order to eliminate the time periods between runs in which no stimulus was present. Channels with a variance smaller than -2 or larger than 2 standard deviations (SD) of the mean activity of all channels were rejected and a notch filter of 50 Hz was applied to remove the line noise.

Independent Component Analysis (ICA) (69) was performed over all remaining channels. Independent components representing eye blinks, eye movements, muscle artifacts, and other types of noise were removed from the EEG signal after which the signals from the previously rejected channels were replaced with the weighted average activity in all remaining channels by spherical spline interpolation. Subsequently, the data were separated into separate datasets containing the report and no-report runs for each participant. The report dataset was segmented into epochs from -2000 to 500 ms around responses. Epochs with response repetitions and epochs that contained more than one response were rejected (22.30% of epochs on average). Finally, epochs were rejected if they exceeded certain thresholds for amplitude (<-150 µV or >150 µV) or slope (>60 µV/epoch), and the data were referenced to the average activity in all channels.

### Eye-tracking recording

Eye movements were recorded at 500 Hz with an EyeLink 1000 (SR Research) infrared eye tracker, calibrated using a 6-point calibration procedure at the start of every block (i.e. five times throughout data collection). During the blocks, participants’ heads were positioned on a chin rest in order to minimize head movements.

### Optokinetic nystagmus (OKN) signal analysis

OKN is a combination of slow-phase and fast-phase eye movements that allows the eyes to follow objects in motion when the head remains stationary, for instance when looking at the trees alongside the road whilst moving past them in a car. The slow-phase movements try to match the stimulus speed to keep the retinal image stable and are interrupted by fast eye movements that reset the eye in orbit. Thus, the velocity of slow-phase OKN provides a continuous and robust estimate of conscious perception rather than the actual visual input, which makes it useful for assessing perception in the absence of reports (70; 25; 28; 71; 26). From the eye movement data, we obtained mean slow-phase OKN velocity and classification accuracy of percepts based on this signal. We performed the following pre-processing steps to obtain the velocity of slow-phase OKN.

First, periods of blinks and saccades were detected using the manufacturer’s standard algorithms with default settings. The subsequent data analyses were performed using custom-made Python software. The following steps were applied to each pupil recording: (i) linear interpolation of values measured just before and after each identified blink (interpolation time window, from 150 ms before until 150 ms after blink), (ii) temporal filtering (third-order Butterworth, low-pass: 10 Hz), (iii) removal of pupil responses to blinks and to saccades, by first estimating these responses by means of deconvolution, and then removing them from the pupil time series by means of multiple linear regression (7), and (iv) conversion to units of modulation (percent signal change) around the mean of the pupil time series from each block.

Second, we smoothed the integrated OKN with a 100 ms Gaussian kernel. We then computed the instantaneous velocity of integrated OKN as the difference between neighboring two-time points (2 ms difference). To obtain the velocity of the slow-phase OKN, we further smoothed the instantaneous velocity with the 100 ms Gaussian kernel. Finally, we segmented the time course of the velocity of slow-phase OKN from 1 s before to 2 s after the onset of stimuli. We did not include the first trial of each block in the analysis as we did not record the fixation position before the first trial.

### Convolutional Neural Network (CNN) classification on button press and OKN

First, to quantify the discriminability of perceptual report in OKN and button press in the report condition, we employed a convolutional neural network (CNN) with the instantaneous velocity of slow-phase OKN as a feature (Figure **??**C,D). The CNN consisted of the following layers in sequence:

A 1D input layer of dimension 1 × 625 units, matching the shape of an epoch of smoothed OKN sampled at 500 Hz.

A convolutional layer consisting of 8 convolutional filters, each with the shape 1 × 25. Units in this layer used ReLU activation functions. The stride step size of the convolution over the inputs was set to 1.

A max pooling layer with the shape 1 × 5, and a stride step size of 2.

A second convolutional layer with 16 convolutional filters, each with the shape 2 × 50. Units used ReLU activation functions and the stride step size of the convolution over the inputs was set to 1.

Another max pooling layer with the shape 1 × 5 and stride step size of 2.

A third (and final) convolutional layer with 32 convolutional filters, each with the shape 2 × 75. Units used ReLU activation functions and the stride step size of the convolution over the inputs was set to 1.

A fully connected dense layer of shape 1 × 2, consisting of units with softmax activation functions.

To prevent overfitting and measure the generalisation performance of the CNN, we first split the OKN data epochs, and their corresponding button press labels into 3 parts: training, validation and testing. 70% of the entire dataset consisting of 13,406 epochs was allocated to the training set. The remaining 30% were further split equally to create validation and test datasets, each containing 15% of the original data and labels. We used stratified sampling when creating these splits, to ensure that the relative proportion of samples of each class was the same in each split.

The CNN was then trained using the training set, with a mini-batch size of 128, over 30 epochs. We randomly shuffled the training dataset at the beginning of each epoch and evaluated the CNN’s performance on the validation dataset once every 10 mini-batches. The stochastic gradient descent algorithm with a momentum of 0.9 was used to learn the weights that minimised the cross-entropy loss over the training dataset. At the end of the training procedure, we recorded the accuracy with which the trained CNN classified OKN epochs in the held-out test dataset. The above procedure – including CNN training, validation, and testing— was repeated, after defining epoch labels based on the OKN crossings in the report condition. Finally, we retrained and evaluated the CNN by defining epoch labels based on the OKN crossings in the no-report condition (Figure 1).

### Directed Information (dir-INFO)

In order to quantify the directed functional connectivity between different EEG signals, we used Directed Information (dir-INFO), also known as Transfer Entropy, an information theoretic measure of Wiener-Granger causality (72; 73; 74). Compared to traditional causality detection methods based on linear models (e.g. Granger causality), dir-INFO is a model-free measure and can detect both linear and nonlinear functional relationships between brain signals. We took advantage of previous work that made this measure statistically robust when applied to neural data (75; 36; 33; 37).

Thus, dir-INFO quantifies functional connectivity by measuring the degree to which the past of a signal *X* predicts the future of another signal *Y*, conditional on the past of *Y*, defined at a specific lag or delay τ: dir-INFO = *I*(*Y*_*t*_; *X*_*t*−τ_ |*Y*_*t*−τ_). Thus, if there is significant dir-INFO between EEG signal *X* at one time, and EEG signal *Y* at a later time, this shows that signal X contains information about the future signal *Y*. Conditioning out the past of signal *Y* ensures the delayed interaction is providing new information over and above that available in the past of signal *X*. For all dir-INFO analyses, we tested delays from 0 ms to 500 ms in steps of 4 ms.

### Information Differentiation (diff-INFO)

We computed Kolmogorov-Chaitin complexity (K complexity) as a metric of information differentiation (76; 77). K complexity quantifies the algorithmic complexity (i.e. the diversity of information patterns) of an EEG signal by measuring its degree of redundancy: from a highly redundant signal (i.e. less differentiated signal) to a slightly redundant one (i.e. highly differentiated signal) (27; 38; 39; 40). Algorithmic complexity of a given EEG sequence can be described as the length of the shortest computer program that can generate it. A short program corresponds to a less complex sequence. K complexity was estimated by quantifying the compression size of the EEG using the Lempel-Ziv zip algorithm (78).

Algorithmic information theory has been introduced by Andreï Kolmogorov and Gregory Chaitin as an area of interaction between computer science and information theory. The concept of algorithmic complexity or Kolmogorov-Chaitin complexity (K complexity) is defined as the shortest description of a string (or in our case a signal *X*). That is to say, K complexity is the size of the smallest algorithm (or computer program) that can produce that particular time series. However, it can be demonstrated by reductio ad absurdum that there is no possible algorithm that can measure K complexity (79). To sidestep this issue, we can estimate an upper-bound value of K complexity(*X*). This can be concretely accomplished by applying a lossless compression of the time series and quantifying the compression size. Capitalizing on the vast signal compression literature, we heuristically used a classical open-source compressor gzip (80) to estimate K complexity(*X*). It is important to standardize the method of representation of the signal before compression in order to avoid irrelevant differences in complexity. Specifically, to compute K complexity(*X*):

First, the signals were transformed into sequences of symbols. Each symbol represents, with identical complexity, the amplitude of the corresponding channel for each time point. The number of symbols was set to 32 and each one corresponds to dividing the amplitude range of that given channel into 32 equivalent bins. Similar results have been obtained with binning ranging from 8 to 128 bins (40). Next, time series were compressed using the compressLib library for Matlab, this library implements the gzip algorithm to compress Matlab variables.

Finally, K complexity(*X*) was calculated as the size of the compressed variable with time series divided by the size of the original variable before compression. Our premise is that the bigger the size of the compressed string, the more complex the structure of the time series, thus indexing the complexity of the electrical activity recorded at an electrode. For each trial and ROI, K complexity was estimated using a 100 ms sliding window with a 4 ms time step.

### EEG time-frequency analysis

Epochs were grouped based on conditions: report, report (OKN), and no report (OKN). Then, EEG traces were decomposed into time-frequency charts from 2 Hz to 26 Hz in 13 linearly spaced steps (2 Hz per bin). The power spectrum of the EEG signal (as obtained by the fast Fourier transform) was multiplied by the power spectra of complex Morlet wavelets !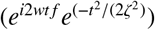 with logarithmically spaced cycle sizes ranging from 3 to 12. The inverse Fourier transform was then used to acquire the complex signal, which was converted to frequency-band specific power by squaring the result of the convolution of the complex and real parts of the signal (*real*[*z*(*t*)]^2^ + *imag*[*z*(*t*)]^2^). The resulting time-frequency data were then averaged per subject and trial type. Finally, time-frequency traces were transformed to decibels (dB) and normalized to the length of the entire trial (baseline: -2000ms to 500 ms) according to: *dB* = 10 ∗ log_10_(power/baseline)

### EEG electrode selection (ROI)

Canonical bilateral frontal (n = 6), parietal (n = 6), as well as right temporal (n = 6) and left temporal (n = 6) electrode clusters were selected for dir-INFO and diff-INFO analyses (Figure S1). Values within each region of interest (ROI) were averaged before computing dir-INFO and diff-INFO metrics and subsequently averaged per condition and participant.

### Statistical analysis

To test for significant time-delay dir-INFO (Figure 2 and 3), and time-frequency spectral power (Figure 5) a cluster-based nonparametric statistical test implemented in FieldTrip (Maris and Oostenveld 2007) was used. In brief, time-delay dir-INFO charts (−2000 to 500 ms) were compared in pairs of experimental conditions (perceptual switches: to INT vs. to DIFF). For each such pairwise comparison, epochs in each condition were averaged subject-wise. These averages were passed to the analysis procedure of FieldTrip, the details of which are described elsewhere (81). In short, this procedure compared corresponding temporal points in the subject-wise averages using dependent samples t-tests for within-subject comparisons. Although this step was parametric, FieldTrip uses a nonparametric clustering method to address the multiple comparisons problem. t values of adjacent temporal points whose p values were lower than 0.01 were clustered together by summating their t values, and the largest such cluster was retained. This whole procedure, i.e., calculation of t values at each temporal point followed by clustering of adjacent t values, was then repeated 1000 times, with recombination and randomized resampling of the subject-wise averages before each repetition. This Monte Carlo method generated a non-parametric estimate of the p-value representing the statistical significance of the originally identified cluster. The cluster-level t value was calculated as the sum of the individual t values at the points within the cluster.

In the case of dir-INFO (Figure 2 and 3, in order to test a possible interaction effect between information direction and perceptual switch, a separate cluster-based permutation test was calculated as the difference between perceptual switch (to INT, to DIFF) and direction of change (front-to-back, back-to-front). Similarly, for diff-INFO analysis, we performed a cluster-based permutation test across the diff-INFO time series (−2000 to 500 ms) between perceptual switches for the front and back ROIs separately. Next, in order to test for an interaction between perceptual switch and direction of change, we performed another cluster-based permutation test on the difference between perceptual switch (to INT, to DIFF) and ROI (front, back).

In case of null findings, we performed a Bayesian RANOVA with identical parameters and settings on the same data, to test if there was actual support of the null hypothesis. When reported, BF_01_ refers to the Bayes Factor in favor of the null hypothesis. Statistical analyses were performed using MATLAB (2019a), Jamovi (Version 0.8.1.6) [Computer Software] (Retrieved from https://www.jamovi.org) (open source), and JASP Team (2018; JASP; version 0.8.4 software) statistical software.

---

**Figure S1:**
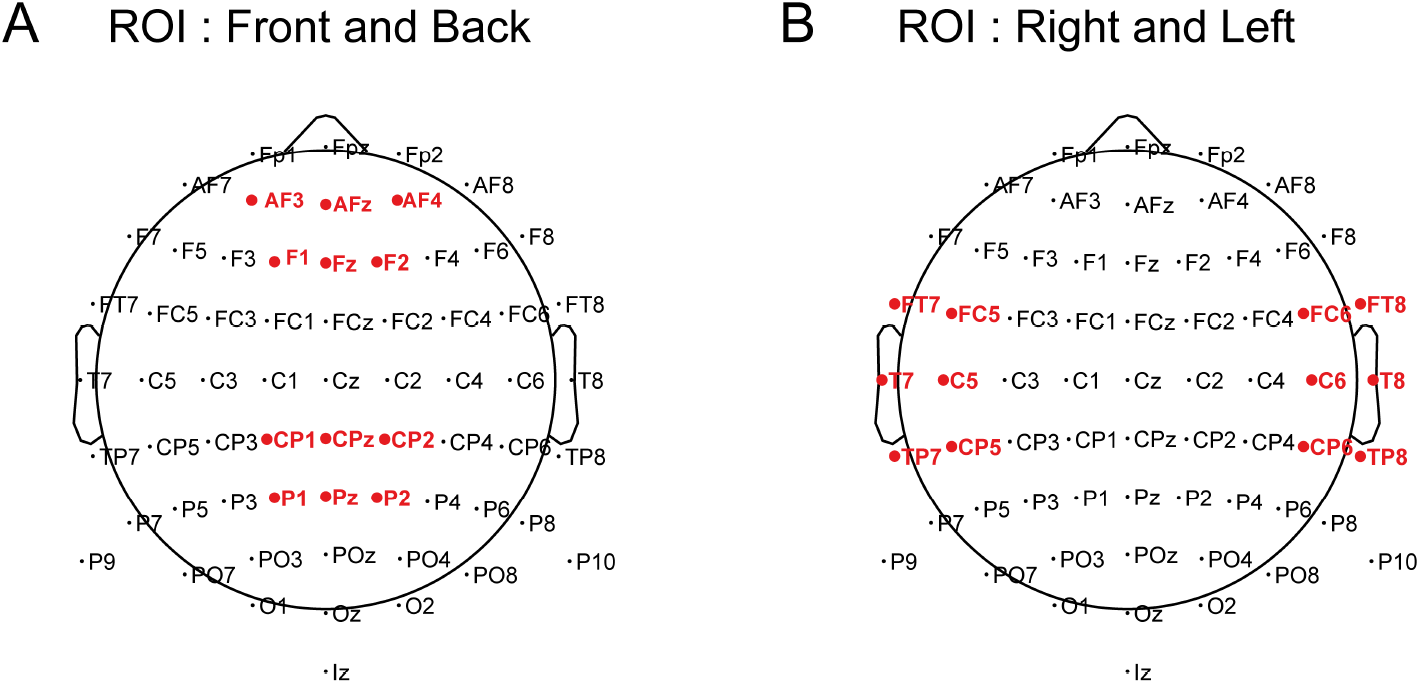
Electrodes included in the **A** Front and Back ROIs, and **B** Right and Left ROIs.

**Figure S2:**
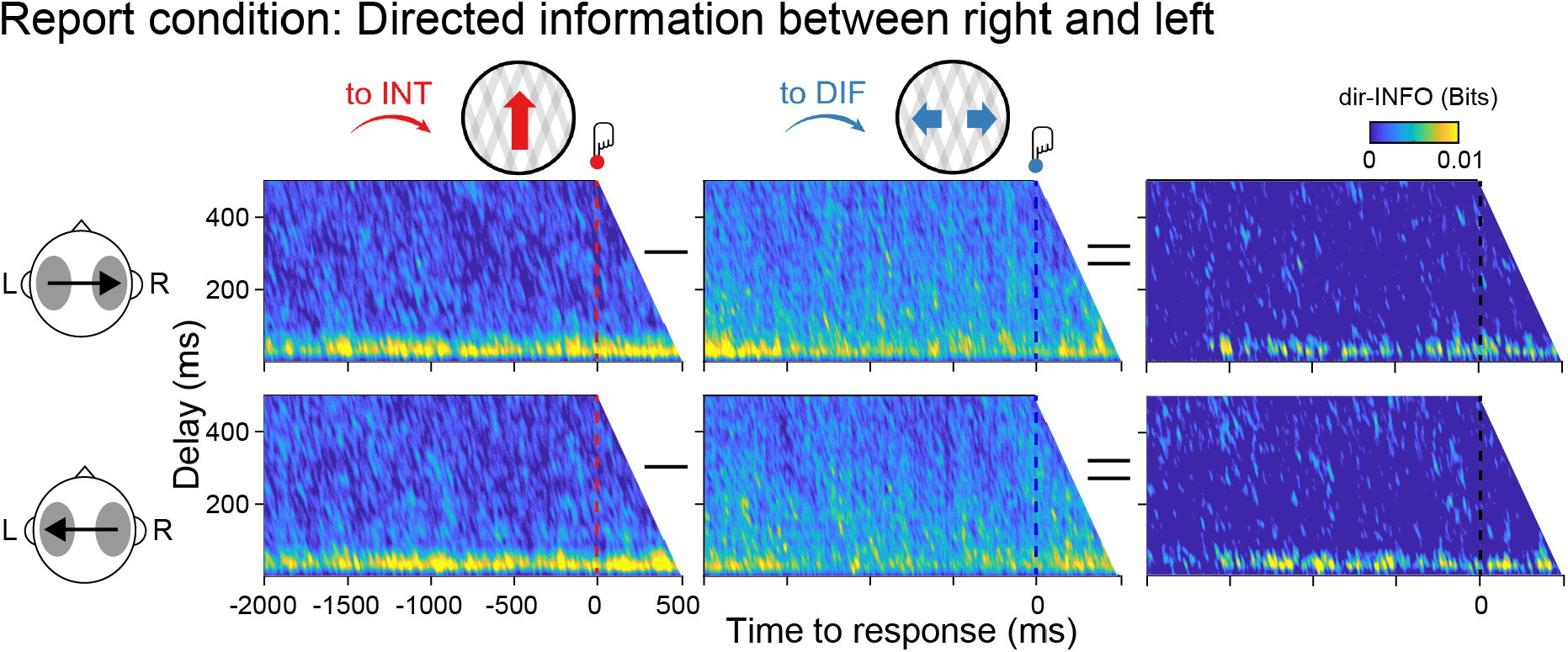
Directed information of the report condition locked to button responses. Group-level dir-INFO between right and left ROIs (upper row) and between left and right ROIs (lower row) when moving plaids are reported as integrated (to INT; red color), reported as differentiated (to DIF; blue color). No significant clusters were found.

**Figure S3:**
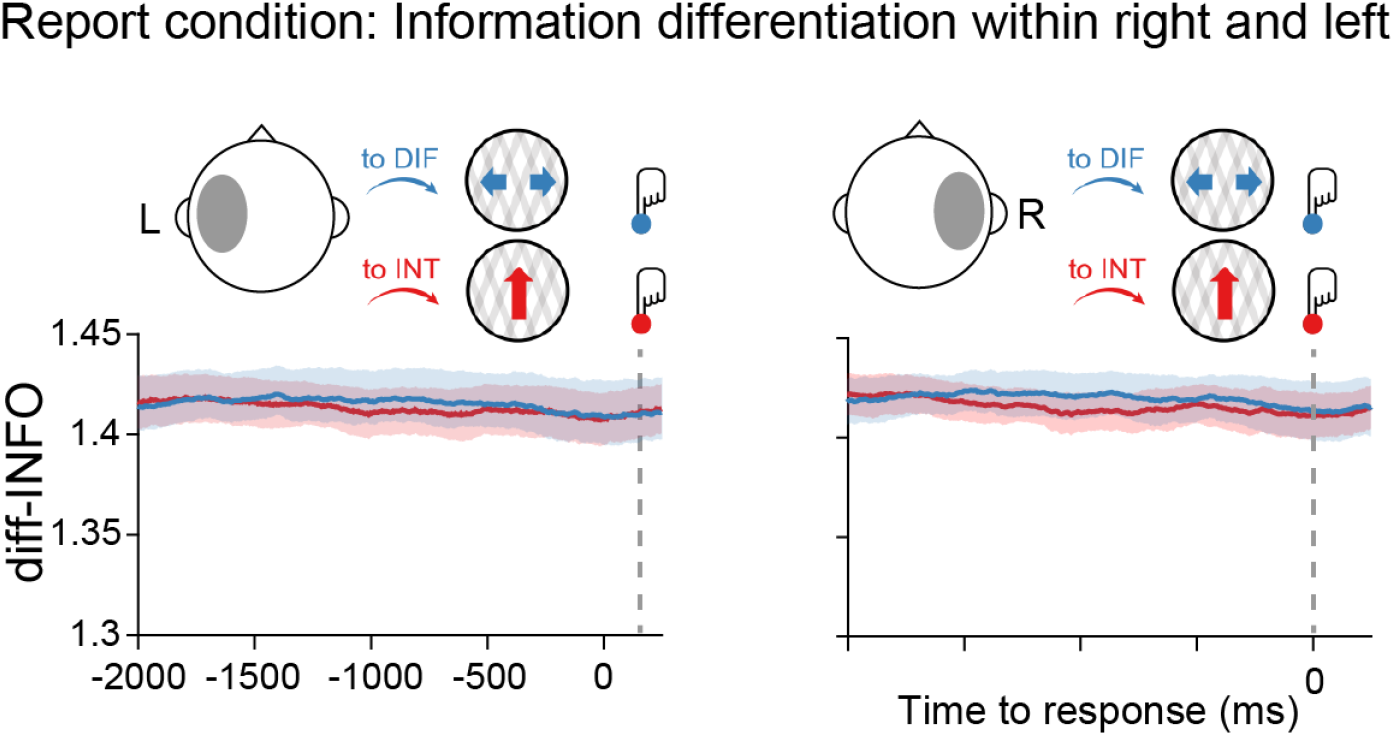
Information differentiation of the report condition locked to button responses. Group-level diff-INFO within the left temporal ROI (left panel) and right temporal (right panel) when moving plaids are reported as differentiated and reported as integrated. No significant clusters were found.

**Figure S4:**
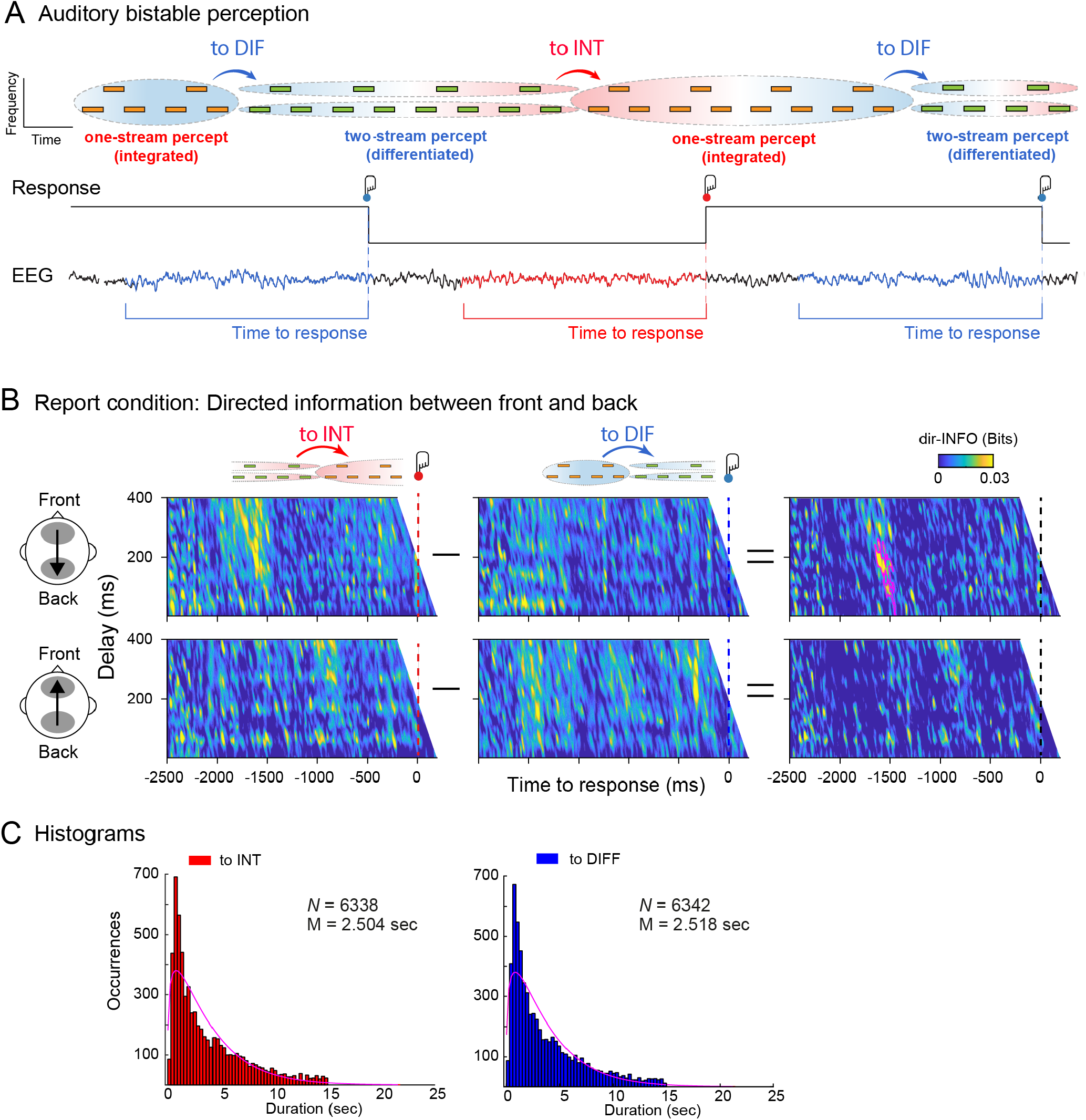
Information dynamics of the report condition locked to button responses. **A** Phenomenology during auditory bistability. Participants (N=29) listened to an ambiguous auditory stream that is experienced either as one auditory stream (integrated percept; red arrow) or two auditory streams (differentiated percept; blue arrows). Perceptual transitions occur either in integrated to differentiated direction (to DIF; blue) or in the differentiated to integrated direction (to INT; red). Middle row: Behavioral responses during the task. Participants pressed one button when perceiving that the integrated percept had fully changed into the differentiated percept (red button) and another button when perceiving that the differentiated percept had fully changed into the integrated percept (blue button). Bottom row: dir-INFO analyses for EEG signals before the button press. **B** Group-level dir-INFO from front to back ROIs (upper row) and from back to front ROIs (lower row) when the auditory stream is reported as integrated (to INT; red color), reported as differentiated (to DIF; blue color), and the cluster-based permutation tests between the two. Note that the significant cluster is observed ∼1500 ms before the response, indicating that a perceptual switch has taken place, which is slightly longer than for our visual bistability results reported in Figure 2A. This minor time difference is likely due to the fact that perceptual states alternated slower in the auditory compared to the visual version of the task. **C** Histograms for perceptual switches in the report condition locked to the button response. These results are re-analyses of the full EEG dataset reported in (27). Experimental design, EEG preprocessing, anterior and posterior electrode selection, and histograms analyses are described in detail in (27)

## Notes

https://osf.io/a2f3v/

### Competing Interest Statement

The authors have declared no competing interest.

### Summary of Updates

This version includes a relevant control analysis performed in an independent dataset (N=29) and reported in the main text and in Supplementary Figure 4.

